# Soft selective sweeps in complex demographic scenarios

**DOI:** 10.1101/004424

**Authors:** Benjamin A. Wilson, Dmitri A. Petrov, Philipp W. Messer

**Affiliations:** Department of Biology, Stanford University, Stanford, CA 94305

**Keywords:** adaptation, mutation, coalescent theory

## Abstract

Recent studies have shown that adaptation from *de novo* mutation often produces so-called soft selective sweeps, where adaptive mutations of independent mutational origin sweep through the population at the same time. Population genetic theory predicts that soft sweeps should be likely if the product of the population size and the mutation rate towards the adaptive allele is sufficiently large, such that multiple adaptive mutations can establish before one has reached fixation; however, it remains unclear how demographic processes affect the probability of observing soft sweeps. Here we extend the theory of soft selective sweeps to realistic demographic scenarios that allow for changes in population size over time. We first show that population bottlenecks can lead to the removal of all but one adaptive lineage from an initially soft selective sweep. The parameter regime under which such ‘hardening’ of soft selective sweeps is likely is determined by a simple heuristic condition. We further develop a generalized analytical framework, based on an extension of the coalescent process, for calculating the probability of soft sweeps under arbitrary demographic scenarios. Two important limits emerge within this analytical framework: In the limit where population size fluctuations are fast compared to the duration of the sweep, the likelihood of soft sweeps is determined by the harmonic mean of the variance effective population size estimated over the duration of the sweep; in the opposing slow fluctuation limit, the likelihood of soft sweeps is determined by the instantaneous variance effective population size at the onset of the sweep. We show that as a consequence of this finding the probability of observing soft sweeps becomes a function of the strength of selection. Specifically, in species with sharply fluctuating population size, strong selection is more likely to produce soft sweeps than weak selection. Our results highlight the importance of accurate demographic estimates over short evolutionary timescales for understanding the population genetics of adaptation from *de novo* mutation.

## Introduction

Adaptation can proceed from standing genetic variation or mutations that are not initially present in the population. When adaptation requires *de novo* mutations, the waiting time until adaptation occurs depends on the product of the mutation rate towards adaptive alleles and the population size. In large populations, or when the mutation rate towards adaptive alleles is high, adaptation can be fast, whereas in small populations the speed of adaptation will often be limited by the availability of adaptive mutations.

Whether adaption is mutation-limited or not has important implications for the population dynamics of adaptive alleles. In a mutation-limited scenario, only a single adaptive mutation typically sweeps through the population and all lineages in a population sample that carry the adaptive allele coalesce into a single ancestor with the adaptive mutation (Figure 1A). This process is referred to as a ‘hard’ selective sweep (Hermisson and Pennings 2005). Hard selective sweeps leave characteristic signatures in population genomic data, such as a reduction in genetic diversity around the adaptive site (Maynard Smith and Haigh 1974; Kaplan *et al.* 1989; Kim and Stephan 2002) and the presence of a single, long haplotype (Hudson *et al.* 1994; Sabeti *et al.* 2002; Voight *et al.* 2006). In non-mutation-limited scenarios, by contrast, several adaptive mutations of independent origin can sweep through the population at the same time, producing so-called ‘soft’ selective sweeps (Pennings and Hermisson 2006a). In a soft sweep, the lineages that carry the adaptive allele collapse into distinct clusters and several haplotypes can be frequent in the population (Figure 1A). As a result, soft sweeps leave more subtle signatures in population genomic data than hard sweeps and are thus more difficult to detect. For example, diversity is not necessarily reduced in the vicinity of the adaptive locus in a soft sweep (Pennings and Hermisson 2006b).

**Figure 1:**
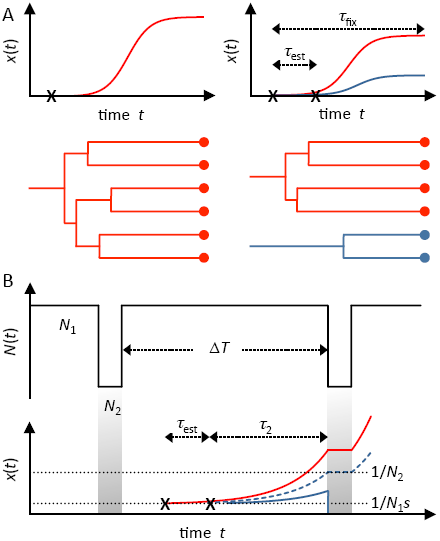
Hard and soft sweeps in populations of constant size and under recurrent population bottlenecks. (A) Allele frequency trajectories and corresponding coalescent genealogies for a hard selective sweep (left) and a soft selective sweep (right). In the soft sweep scenario, a second beneficial mutation establishes *τ*_est_ generations after the first mutation but before the beneficial allele has fixed. The distinguishing feature between a hard and a soft sweep can be seen in the genealogy of a population sample of individuals with the adaptive allele: in a hard sweep, the sample coalesces into a single ancestor, whereas in a soft sweep the sample coalesces into multiple ancestors with independently arisen adaptive mutations. (B) Illustration of our simplified model used to explore the hardening phenomenon. Population bottlenecks occur every Δ*T* generations wherein the population size is reduced from *N*_1_ to *N*_2_ for a single generation. The average waiting time between independently establishing beneficial mutations is *τ*_est_. From establishment, it takes *τ*_2_ generations for the second mutation to reach frequency 1/*N*_2_, from where on it is unlikely to be lost during the bottleneck. The hardening phenomenon is illustrated by the loss of the dark blue allele during the bottleneck. The dashed blue line indicates the threshold trajectory required for the mutation to successfully survive the bottleneck.

There is mounting evidence that adaptation is not mutation-limited in many species, even when it requires a specific nucleotide mutation in the genome (Messer and Petrov 2013). Recent case studies have revealed many examples where, at the same locus, several adaptive mutations of independent mutational origin swept through the population at the same time, producing soft selective sweeps. For instance, soft sweeps have been observed during the evolution of drug resistance in HIV (Fischer *et al.* 2010; Messer and Neher 2012; Pennings *et al.* 2014) and malaria (Nair *et al.* 2007), pesticide and viral resistance in fruit flies (Catania *et al.* 2004; Aminetzach *et al.* 2005; Chung *et al.* 2007; Karasov *et al.* 2010; Schmidt *et al.* 2010), warfarin resistance in rats (Pelz *et al.* 2005), and color patterns in beach mice (Hoekstra *et al.* 2006; Domingues *et al.* 2012). Even in the global human population, adaptation has produced soft selective sweeps, as evidenced by the parallel evolution of lactase persistence in Eurasia and Africa through recurrent mutations in the lactase enhancer (Bersaglieri *et al.* 2004; Tishkoff *et al.* 2007; Enattah *et al.* 2008; Jones *et al.* 2013) and the mutations in the gene *G6PD* that evolved independently in response to malaria (Louicharoen *et al.* 2009). Some of these sweeps arose from standing genetic variation while others involved recurrent *de novo* mutation. For the remainder of our study, we will focus on the latter scenario of adaptation arising from *de novo* mutation.

The population genetics of adaptation by soft selective sweeps was first investigated in a series of papers by Hermisson and Pennings (Hermisson and Pennings 2005; Pennings and Hermisson 2006a,b). They found that in a haploid population of constant size the key evolutionary parameter that determines whether adaptation from *de novo* mutations is more likely to produce hard or soft sweeps is the population-scale mutation rate Θ = 2*N*_*e*_*U*_*A*_, where *N*_*e*_ is the variance effective population size in a Wright-Fisher model and *U*_*A*_ is the rate at which the adaptive allele arises per individual per generation. When Θ ≪ 1, adaptation typically involves only a single adaptive mutation and produces a hard sweep, whereas when Θ becomes on the order of one or larger, soft sweeps predominate (Pennings and Hermisson 2006a).

The strong dependence of the likelihood of soft sweeps on Θ can be understood from an analysis of the involved timescales. An adaptive mutation with selection coefficient *s* that successfully escapes early stochastic loss requires *τ*_fix_ ≈ log(*N*_*e*_*s*)/*s* generations until it eventually fixes in the population (Hermisson and Pennings 2005; Desai and Fisher 2007). The expected number of independent adaptive mutations that arise during this time is on the order of *NU*_*A*_ log(*N*_*e*_*s*)/*s* – *i.e.*, the product of the population-scale mutation rate towards the adaptive allele and its fixation time. Yet only an approximate fraction 2*s* of these mutations will escape early stochastic loss and successfully establish in the population (Haldane 1927; Kimura 1962). Thus, the expected number of independently originated adaptive mutations that successfully establish before the first one has reached fixation is of order (2*s*)*NU*_*A*_ log(*N*_*e*_*s*)/*s* = Θ log(*N*_*e*_*s*) and, therefore, depends only logarithmically on the selection coefficient of the adaptive allele.

Our current understanding of the likelihood of soft sweeps relies on the assumption of a Wright-Fisher model with fixed population size, where Θ remains constant over time. This assumption is clearly violated in many species, given that population sizes often change dramatically throughout the evolutionary history of a species. In order to assess what type of sweeps to expect in a realistic population, we must understand how the likelihood of soft sweeps is affected by demographic processes.

In many organisms population sizes can fluctuate continuously and over timescales that are not necessarily long compared to those over which adaptation occurs. For example, many pathogens undergo severe bottlenecks during host-to-host transmission (Artenstein and Miller 1966; Gerone *et al.* 1966; Wolfs *et al.* 1992; Wang *et al.* 2010), insects can experience extreme, seasonal boom-bust cycles (Wright *et al.* 1942; Ives 1970; Baltensweiler and Fischlin 1988; Nelson *et al.* 2013), and even some mammals experience dramatic, cyclical changes in abundance (Myers 1998; Krebs and Myers 1974). Extensive work has been devoted to the question of how such fluctuations affect the fixation probabilities of adaptive mutations (Ewens 1967; Otto and Whitlock 1997; Pollak 2000; Patwa and Wahl 2008; Engen *et al.* 2009; Parsons *et al.* 2010; Uecker and Hermisson 2011; Waxman 2011) but it remains unclear how they affect the likelihood of observing soft sweeps.

In this study we investigate the effects of demographic processes on adaptation from *de novo* mutations. We show that recurrent population bottlenecks can give rise to a phenomenon we term the ‘hardening’ of soft selective sweeps. Hardening occurs when only one beneficial lineage in an initially soft sweep persists through a population bottleneck. We then develop a generalized analytical framework for calculating the likelihood of soft sweeps under arbitrary demographic scenarios, based on the coalescent with ‘killings’ process. We find that when population size varies over time, two important symmetries of the constant population size scenario are broken: first, the probability of observing soft sweeps becomes a function of the starting time of the sweep and, second, it becomes a function of the strength of selection. In particular, we show that strong selection is often more likely to produce soft sweeps than weak selection when population size fluctuations are common.

## Results

We study a single locus with two alleles, *a* and *A*, in a haploid Wright-Fisher population (random mating, discrete generations) (Ewens 2004). The population is initially monomorphic for the wildtype allele *a*. The derived allele *A* has a selective advantage *s* over the wildtype and arises at a rate *U*_*A*_ per individual, per generation. We ignore back mutations and consider the population dynamics of the two alleles at this locus in isolation, *i.e.*, there is no interaction with other alleles elsewhere in the genome.

In a classical hard sweep scenario, a single adaptive allele arises, successfully escapes early stochastic loss, and ultimately sweeps to fixation in the population. In a soft sweep, several adaptive mutations establish independently in the population and rise in frequency before the adaptive allele has fixed in the population. After fixation of the adaptive allele, the lineages in a population sample do not coalesce into a single ancestor with the adaptive allele but fall into two or more clusters, reflecting the independent mutational origins of the different adaptive lineages (Figure 1A). Note that the distinction between a hard and a soft sweep is based on the genealogy of adaptive alleles in a population sample. It is therefore possible that the same adaptive event yields a soft sweep in one sample but remains hard in another, depending on which individuals are sampled.

### Soft sweeps in populations of constant size

The likelihood of soft sweeps during adaptation from *de novo* mutation has been calculated by Pennings and Hermisson (2006a) for a Wright-Fisher model of constant population size *N*. Using coalescent theory, they showed that in a population sample of size *n*, drawn right after fixation of the adaptive allele, the probability of observing at least two independently originated adaptive lineages is given by

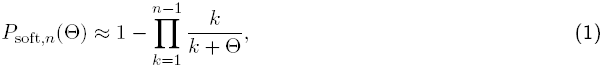

where Θ = 2*NU*_*A*_ is the population-scale mutation rate – twice the number of adaptive lineages that enter the population per generation. Thus, the probability of a soft sweep is primarily determined by Θ and is nearly independent of the strength of selection.

The transition between the regimes where hard and where soft sweeps predominate occurs when Θ becomes on the order of one in the constant population size scenario. When Θ ≪ 1, adaptive mutations are not readily available in the population and adaptation is impeded by the waiting time until the first successful adaptive mutation arises. This regime is referred to as the mutation-limited regime. Adaptation from *de novo* mutation typically produces hard sweeps in this case. When Θ ≥ 1, by contrast, adaptive mutations arise at least once per generation on average. In this non-mutation-limited regime, soft sweep predominate.

### Soft sweeps under recurrent bottlenecks: heuristic predictions

The standard Wright-Fisher model assumes a population of constant size *N*. To study the effects of population size changes on the probability of soft sweeps, we relax this condition and model a population that alternates between two sizes. Every Δ*T* generations the population size is reduced from *N*_1_ to *N*_2_ ≪ *N*_1_ for a single generation and then returns to its initial size in the following generation (Figure 1B). We define Θ = 2*N*_1_*U*_*A*_ as the population-scale mutation rate during the large population phases.

We assume instantaneous population size changes and do not explicitly consider a continuous population decline at the beginning of the bottleneck or growth during the recovery phase. This assumption should be appropriate for sharp, punctuated bottlenecks and allows us to specify the ‘severity’ of a bottleneck in terms of a single parameter, *N*_2_/*N*_1_. Note that many effects of a population bottleneck depend primarily on the ratio of its duration over its severity. In principle, most of the results we derive below should therefore be readily applicable to more complex bottleneck scenarios by mapping the real bottleneck onto an effective single-generation bottleneck. We also assume that mutation and selection are only operating during the phases when the population is large, whereas the two alleles, *a* and *A*, are neutral with respect to each other and no new mutations occur during a bottleneck. This assumption is justified for severe bottlenecks with *N*_2_ ≪ *N*_1_.

Adaptive mutations arise in the large population at rate *N*_1_*U*_*A*_, but only a fraction 2*s* of these mutations successfully establishes in the large population, *i.e.*, these mutations stochastically reach a frequency ≈ 1/(*N*_1_*s*) whereupon they are no longer likely to become lost by random genetic drift (assuming that the amount of drift remains constant over time). Thus, adaptive mutations establish during the large phases at an approximate rate Θ_*s*_. We assume that successfully establishing mutations reach their establishment frequency fast compared to the timescale Δ*T* between bottlenecks, in which case establishment can be effectively modeled by a Poisson process. This assumption is reasonable when selection is strong and the establishment frequency low. Note that those adaptive mutations that do reach establishment frequency typically achieve this within very few generations (Eriksson *et al.* 2008).

Under the Poisson assumption, the expected waiting time until an adaptive mutation successfully establishes in the large population phase is given by *τ*_est_ = 1/(Θ*s*). After establishment, its population frequency is modeled deterministically by logistic growth: *x*(*t*) = 1/[1 + (*N*_1_*s*) exp(−*st*)]. Fixation would thus occur within *τ*_fix_ ≈ log(*N*_1_*s*)/*s* generations after establishment, assuming that the population size were to remain constant.

If an adaptive mutation establishes during the large phase but has not yet fixed at the time the next bottleneck occurs, its fate will depend on its frequency at the onset of the bottleneck. In our model, the bottleneck is a single generation of random down-sampling of the population to a size *N*_2_ ≪ *N*_1_. Any allele present at the onset of the bottleneck will likely survive the bottleneck only when it was previously present at a frequency larger than 1/*N*_2_, *i.e.*, when at least one copy of the allele is expected to be present during the bottleneck. Less frequent alleles will typically be lost (Figure 1B). To reach frequency 1/*N*_2_ in the population, an adaptive mutation needs to grow for approximately another *τ*_2_ = log(*N*_1_*s*/*N*_2_)/*s* generations after establishment. We can therefore define the ‘bottleneck establishment time’ as the sum of the initial establishment time, *τ*_est_, and the waiting time until the mutation has subsequently reached a high-enough frequency to likely survive a bottleneck, *τ*_2_:

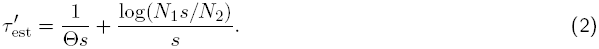

We will show below that the comparison between bottleneck establishment time, 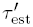, and bottleneck recurrence time, Δ*T*, distinguishes the qualitatively different regimes in our model.

**Mutation-limited adaptation:** It is clear that bottlenecks can only decrease the probability of a soft sweep in our model relative to the probability in the constant population size scenario, as they systematically remove variation from the population by increasing the variance in allele frequencies between generations. Consequently, when Θ ≪ 1 sweeps will be hard because adaptation is already mutation-limited during the large phases. Note that mutation-limitation does not necessarily imply that adaptation is unlikely in general, it may just take longer until an adaptive mutation successfully establishes in the population. When the recurrence time, Δ*T*, is much larger than the establishment time, *τ*_est_, adaptation is still expected to occur between two bottlenecks.

**Non-mutation-limited adaptation:** If Θ ≥ 1, adaptation is not mutation-limited during the large population phases. In the absence of bottlenecks (or when bottlenecks are very weak), adaptation from *de novo* mutation will often produce soft selective sweeps. A strong population bottleneck, however, can potentially remove all but one adaptive lineage and result in a scenario where only this one lineage ultimately fixes. In this case, we say that the bottleneck has ‘hardened’ the initially soft selective sweep.

We can identify the conditions that make hardening likely from a simple comparison of timescales: hardening should occur whenever Θ ≥ 1 and at the same time

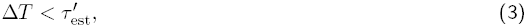

such that a second *de novo* mutation typically does not have enough time to reach a safe frequency that assures its survival before the next bottleneck sets in (Figure 1B).

The argument that the second adaptive mutation needs to grow for *τ*_2_ generations after its establishment to reach a safe frequency 1/*N*_2_ only makes sense when the mutation is actually at a lower frequency than 1/*N*_2_ at establishment, which requires that bottlenecks are sufficiently severe (*N*_2_/*N*_1_ < *s*). For weaker bottlenecks, most established mutations should typically survive the bottleneck and hardening will generally be unlikely. Note that the condition *N*_2_/*N*_1_ > s alone does not imply that soft sweeps should predominate – this still depends on the value of Θ. In the other limit, where bottleneck severity increases until *N*_2_ → 1, all sweeps become hardened. This imposes the requirement that *τ*_2_ ≪ *τ*_fix_ or correspondingly that *N*_2_ ≫ 1 for our bottleneck establishment time to be valid.

The heuristic argument invokes a number of strong simplifications, including that allele frequency trajectories are deterministic once the adaptive allele has reached its establishment frequency, that alleles at frequencies below 1*/N*_2_ have no chance of surviving a bottleneck, and that establishment occurs instantaneously during a large population phase. In reality, however, an adaptive mutation spends time in the population before establishment. And if this time becomes on the order of Δ*T*, then adaptive mutations encounter bottlenecks during the process of establishment. In this case, establishment frequency will be higher than 1/(*N*_1_*s*) and establishment time will be longer than 1/(Θ*s*) due to the increased drift during bottlenecks. We will address these issues more thoroughly below when we analyze general demographic scenarios.

Our condition relating the bottleneck recurrence time and the bottleneck establishment time (3) makes the interesting prediction that for fixed values of Θ, Δ*T*, and *N*_2_*/N*_1_, there should be a threshold selection strength for hardening. Sweeps involving weaker selection than this threshold are likely to be hardened, whereas stronger sweeps are not. Thus, both hard and soft sweeps can occur in the same demographic scenario, depending on the strength of selection. This is in stark contrast to the constant population size scenario, where primarily the value of Θ determines whether adaptation produces hard or soft sweeps while the strength of selection enters only logarithmically.

### Soft sweeps under recurrent bottlenecks: forward simulations

We performed extensive forward simulations of adaptation from *de novo* mutation under recurrent population bottlenecks to measure the likelihood of soft sweeps in our model and to assess the accuracy of condition (3) under a broad range of parameter values. In our simulations we modeled the dynamics of adaptive lineages at a single locus in a modified Wright-Fisher model with selection (Methods). To estimate the empirical probability of observing a soft sweep in a given simulation run, we calculated the probability that two randomly sampled adaptive lineages are not identical by decent at the time of fixation of the adaptive allele, *i.e.*, arose from independent mutational origins.

Figure 2 shows phase diagrams of the empirical probabilities of soft sweeps in our simulations over a wide range of parameter values. We investigated three Θ-regimes that differ in the relative proportions at which hard and soft sweeps arise during the large phases before they experience a bottleneck: (i) mostly hard sweeps arise during the large phase (Θ = 0.2), (ii) mostly soft sweeps arise during the large phase (Θ = 2), and (iii) practically only soft sweeps arise during the large phase (Θ = 20). For each value of Θ, we investigated three different bottleneck severities: *N*_1_/*N*_2_ = 10^2^, *N*_2_/*N*_1_ = 10^3^, and *N*_1_/*N*_2_ = 10^4^.

**Figure 2:**
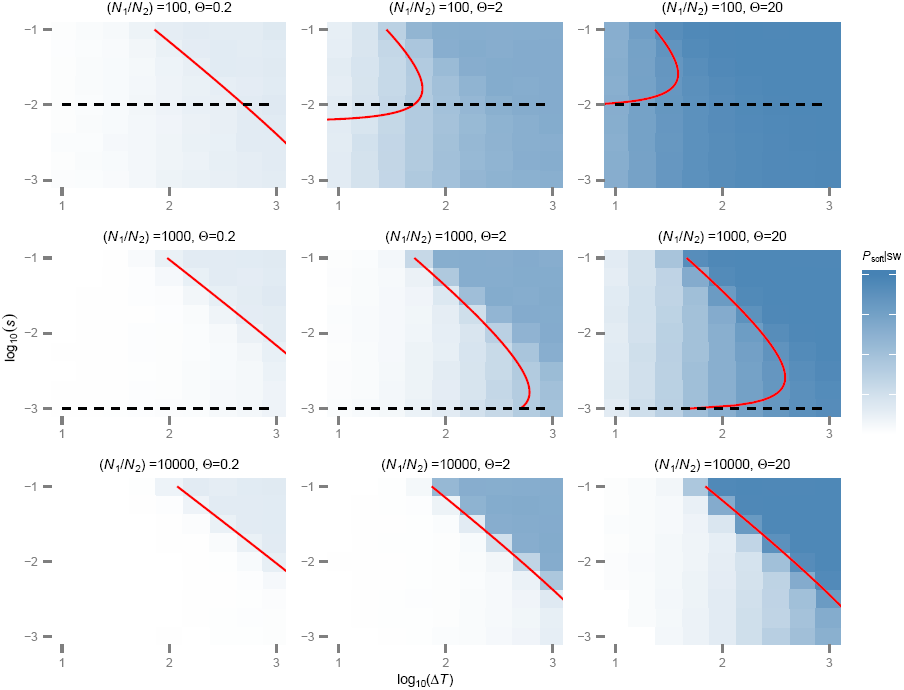
Hardening of soft selective sweeps under recurrent population bottlenecks. The nine panels show different bottleneck severities (weaker to stronger from top to bottom) and different population-scale mutation rates, Θ, during the large population phases. The coloring of the squares specifies the proportion of soft sweeps observed in samples of two individuals at the time of fixation for 1000 simulations runs (Methods) with selection coefficient (*s*) and bottleneck recurrence time (Δ*T*) at the center of each square. The red lines indicate the boundary condition 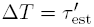 between the regime where hardening is predicted to be likely (left of line) and unlikely (right of line) according to our heuristic Equation (3). The dashed black line indicates the boundary condition *N*_2_/*N*_1_ = *s* on the severity of the bottleneck; below the line, bottlenecks are not severe enough for the hardening condition to be applicable. Note that for the low population-scale mutation rate Θ = 0.2 in the left panels only very few sweeps are soft initially during the large population phase, and hardening therefore is unlikely from the outset.

Our simulations confirm that hardening is common in populations that experience sharp, recurrent bottlenecks. The evolutionary parameters under which hardening is likely are qualitatively distinguished by the heuristic condition (3). Hardening becomes more likely with increasing severity of the population bottlenecks. For a fixed value of Θ and a fixed severity of the bottlenecks, hardening also becomes more likely the weaker the strength of positive selection and the shorter the recurrence time between bottlenecks, as predicted. For the scenarios with Θ = 0.2, most sweeps are already hard when they arise. Thus, there are only few soft sweeps that could be subject to hardening, leading to systematically lower values of *P*_soft_ compared to the scenarios with higher values of Θ. Note that the transition between the regimes where hardening is common and where it is uncommon can be quite abrupt. For example, in the scenario where Θ = 2, *N*_1_/*N*_2_ = 10^4^, and Δ*T* = 100 generations, an adaptive allele with *s* = 0.056 almost always (90%) produced a hard sweep in our simulations, whereas an allele with *s* = 0.1 mostly (57%) produced a soft sweep.

### Probability of soft sweeps in complex demographic scenarios

In this section we describe an approach for calculating the probability of observing soft sweeps from recurrent *de novo* mutation that can be applied to complex demographic scenarios. We assume that the population is initially monomorphic for the wildtype allele, *a*, and that the adaptive allele, *A*, has selection coefficient *s* and arises through mutation of the wildtype allele at rate *U*_*A*_ per individual, per generation. Let *P*_soft,*n*_(*t*_0_, *s*) denote the probability that a sweep arising at time *t*_0_ is soft in a sample of *n* adaptive alleles. Generally *P*_soft, *n*_(*t*_0_, *s*) will also be a function of the trajectory, *x*(*t* ≥ *t*_0_), of the adaptive allele, the specific demographic scenario, *N* (*t* ≥ *t*_0_), and the sampling time, *t*_*n*_.

We can calculate *P*_soft, *n*_(*t*_0_, *s*) given *x*(*t*), *N*(*t*), and *t*_*n*_ using a straightforward extension of the approach employed by Pennings and Hermisson (2006a) in deriving *P*_soft,*n*_(Θ) for a population of constant size, which resulted in Equation (1). In particular, we can model the genealogy of adaptive alleles in a population sample by a coalescent process with ‘killings’ (Durrett 2008). In this process, two different types of events can occur in the genealogy of adaptive alleles when going backwards in time from the point of sampling: two lineages can coalesce, or a lineage can mutate from the wildtype allele to the adaptive allele (Figure 3). In the latter case, the lineage in which the mutation occurred is stopped (referred to as killing). Thus, each pairwise coalescence event and each mutation event reduce the number of ancestral lineages in the genealogy by one. The process stops when the last ancestral lineage is stopped by a mutation (which cannot occur further back in the past than time *t*_0_, the time when the adaptive allele first arose in the population).

**Figure 3:**
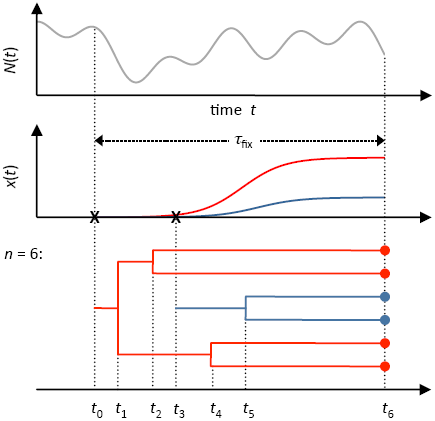
Modeling the genealogy of adaptive alleles by a coalescent process with killings. Population size *N* (*t*) can vary arbitrarily over time in our model (top panel). An adaptive allele arises in the population (indicated by **x**) in generation *t*_0_ and subsequently sweeps through the population (red frequency trajectory *x*(*t*) in middle panel). Before fixation, a second adaptive lineage arises by mutation (indicated by second **x**) and also sweeps through the population (blue frequency trajectory in middle panel). The bottom panel illustrates a possible genealogy of a population sample of *n* = 6 adaptive alleles, taken at the time *t*_6_. When tracing the lineages back in time, a pair of lineages can coalesce (events *t*_1_, *t*_2_, *t*_4_, and *t*_5_) or a lineage can mutate (events *t*_0_ and *t*_3_), indicating *de novo* mutational origin of the adaptive allele. In the latter case the lineage is killed. The shown example is a soft sweep because a second *de novo* mutation occurs before all lineages have coalesced into a single ancestor.

Hard and soft sweeps have straightforward interpretations in this framework: In a hard sweep, all lineages in the sample carry the adaptive allele from the same mutational origin and therefore coalesce into a single ancestral lineage before the process finally stops. In a soft sweep, on the other hand, at least one additional mutation occurs before the process stops (Figure 3).

We will depart from the Wright-Fisher framework here and instead model this coalescent as a continuous-time Markov process. The instantaneous rates of coalescence (*λ*_coal_) and mutation (*λ*_mut_) at time *t*, assuming that *k* ancestral lineages are present in the genealogy at this time, are then given by

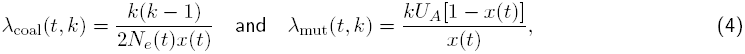

where *N*_*e*_(*t*) is the single-generation variance effective population size in generation *t*. Note that these are the same rates as used by Pennings and Hermisson (2006a), with the only difference being that in our case the population size *N*_*e*_(*t*) can vary over time.

Let us for now assume we were to actually know the times *t*_1_, …, *t*_*n*−1_ at which coalescence or mutation events happen in the genealogy, where *t*_*k*_ for *k* = 1, …, *n* − 1 specifies the time at which the coalescence or mutation event happens that reduces the number of ancestral lineages from *k* + 1 to *k*, and *t*_*n*_ specifies the time of sampling (Figure 3). Note that we do not make any assumptions about when the sample is taken, we only require that there are *n* copies of the adaptive allele present in the sample. Given a pair of successive time points, *t*_*k*_ and *t*_*k*+1_, we can calculate the probability *P*_coal_(*t*_*k*_) that this event is a coalescence event, rather than a mutation event, using the theory of competing Poisson processes:

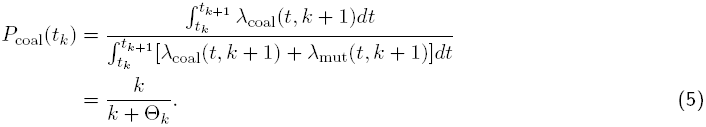

The last equation holds if we define an effective Θ_*k*_ as

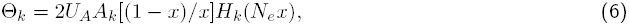

where 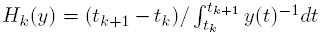 denotes the harmonic mean and 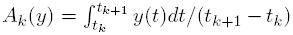 the arithmetic mean, estimated over the interval [*t*_*k*_, *t*_*k*+1_]. This effective Θ_*k*_ recovers the original result Θ_*k*_ = 2*N*_*e*_*U*_*A*_ from Pennings and Hermisson (2006a) for the special case of constant population size, where 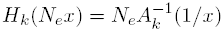 and mutation and coalescence should only be likely during the early phase of a sweep, when *A*_*k*_[(1 − *x*)/*x*] ≈ *A*_*k*_(1/*x*).

The effective Θ_*k*_ from Equation (6) describes the product of two specific means estimated during the time interval between events *k* and *k* + 1: (i) the arithmetic mean of twice the rate at which mutations towards the adaptive allele occur per lineage and (ii) the harmonic mean of *N*_*e*_*x*, the effective number of individuals that carry the adaptive allele at time *t*. The first mean is independent of demography and will be largest during the early phase of a sweep when *x*(*t*) is small. The second mean depends on the product of both the trajectory, *x*(*t*), and the demography, *N*_*e*_(*t*). Importantly, as a harmonic mean, it is dominated by the smallest values of *N*_*e*_*x* during the estimation interval. Thus, even if the estimation interval lies in a later stage of the sweep, when *x*(*t*) is larger than it was early in the sweep, the harmonic mean could nevertheless be small if *N*_*e*_(*t*) is small at some point during this interval. In general, when population size varies over time, it is not always true that most coalescence occurs during the early phase of a sweep, and we will therefore not adopt this assumption here. For instance, if a strong bottleneck is encountered late during the sweep, most coalescence can occur within this bottleneck.

Given an arbitrary demographic scenario, *N*_*e*_(*t*), and trajectory *x*(*t*) of the adaptive allele, Equation (6) allows us to calculate each effective Θ_*k*_ if we know the time points *t*_*k*_ and *t*_*k*+1_. Given the sequence {Θ_*k*_} for all *k* = 1, …, *n* − 1, we can then calculate the probability that the sweep in our sample is hard, as this is only the case if all individual events in the genealogy happen to be coalescence events. The probability that this happens is the product of all *P*_coal_(*t*_*k*_). Hence, the probability that the sweep is soft in our sample is

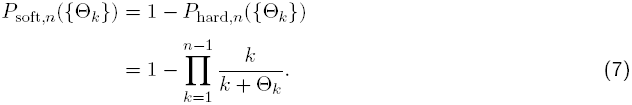

### Calculating Θ_*k*_ for a given demographic scenario

The above calculation of *P*_soft,*n*_ based on Equations (6) and (7) presupposed that we actually know the trajectory of the adaptive allele and the times *t*_*k*_ at which coalescence or mutation events occur in the genealogy.

This assumption is unrealistic in practice. A full treatment of the problem in the absence of such information then requires integrating over all possible trajectories and all individual times at which coalescence or mutation events can occur, where we weigh each particular path *x*(*t*) and sequence of event times *t*_1_, …, *t*_*n*_ by their expected probabilities.

Instead of performing such a complicated ensemble average, we use a deterministic approximation for the trajectory *x*(*t*) and then model the times *t*_*k*_ as stochastic random variables that we approximate by their expectation values. Specifically, we model the frequency trajectory of an adaptive allele in the population by

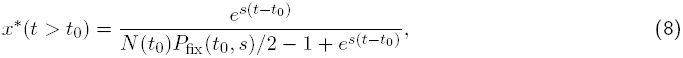

where *P*_fix_(*t*_0_, *s*) is the fixation probability of a new mutation of selection coefficient *s* that arises in the population at time *t*_0_ in a single copy (Uecker and Hermisson 2011). Calculating such fixation probabilities when population size varies over time has been the subject of several studies and is well understood (Ewens 1967; Otto and Whitlock 1997; Pollak 2000; Patwa and Wahl 2008; Engen *et al.* 2009; Parsons *et al.* 2010; Uecker and Hermisson 2011; Waxman 2011). For example, Uecker and Hermisson (2011) have derived the following general formula for calculating *P*_fix_(*t*_0_, *s*) under arbitrary demographic scenarios:

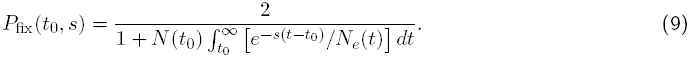

Here *N*_*e*_(*t*) again specifies the single-generation variance effective population size in generation *t*. This approximation works well as long as the number of beneficial mutations that enter the population during the sweep is not extremely high (Θ ≫ 1), in which case one would need to explicitly include the contribution from mutation in the formulation of the birth-death process.

Assuming that the adaptive allele follows the deterministic trajectory, *x*\*(*t*), from Equation (8), we can calculate the expected rates of coalescence, 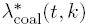, and mutation, 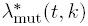, in the genealogy of adaptive alleles in a population sample. Let us assume the sample of size *n* is taken at *t*_*n*_. The expectation value *E*(*t*_*k*_) of the times *t*_*k*_ (*k* = 1, …, *n* − 1) at which the number of lineages goes from *k* + 1 to *k* then obeys the relation

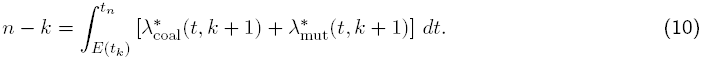

In other words, *E*(*t*_*k*_) is the average waiting time until *n* − *k* events have occurred in the genealogy of the sample when going backwards in time from the point of sampling. Given the expected times *E*(*t*_*k*_), one can then calculate the expected values *E*(Θ_*k*_) via Equation (6) and estimate *P*_soft,*n*_(*t*_0_, *s*) via Equation (7).

### Application for cycling populations

To illustrate and verify our approach for calculating *P*_soft,*n*_(*t*_0_, *s*), we examine selective sweeps in a population that undergoes cyclical population size changes. In particular, we model a haploid Wright-Fisher population with a time-dependent population size given by:

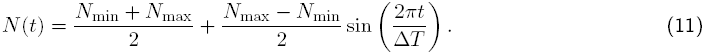

As illustrated in Figure 4A, this specifies a population that cycles between a minimal size, *N*_min_, and a maximal size, *N*_max_, over a period of Δ*T* generations. We investigate selective sweeps with four different starting times (*t*_0_) at which the successfully sweeping allele first arises within a cycle: *t*_0_ = 0, *t*_0_ = 0.25Δ*T*, *t*_0_ = 0.5Δ*T*, and *t*_0_ = 0.75Δ*T*. These four cases describe, in order, a starting time of the sweep midway during a growth phase, at the end of a growth phase, midway of a decline phase, and at the end of a decline phase (Figure 4A). For each starting time we calculate the expected probability *P*_soft_,_2_(*t*_0_, *s*) of observing a soft sweep in a sample of size two as a function of the selection coefficient (*s*, of the adaptive allele, assuming that the population is sampled when the adaptive allele has reached population frequency *x* = 1/2. In contrast to sampling at the time of fixation, this criterion does not depend on the actual population size (*e.g.* in a growing population fixation can take very long). Note that the probability *P*_soft,2_(*t*_0_, *s*) is the probability that two adaptive alleles in a random population sample are not identical by decent.

**Figure 4:**
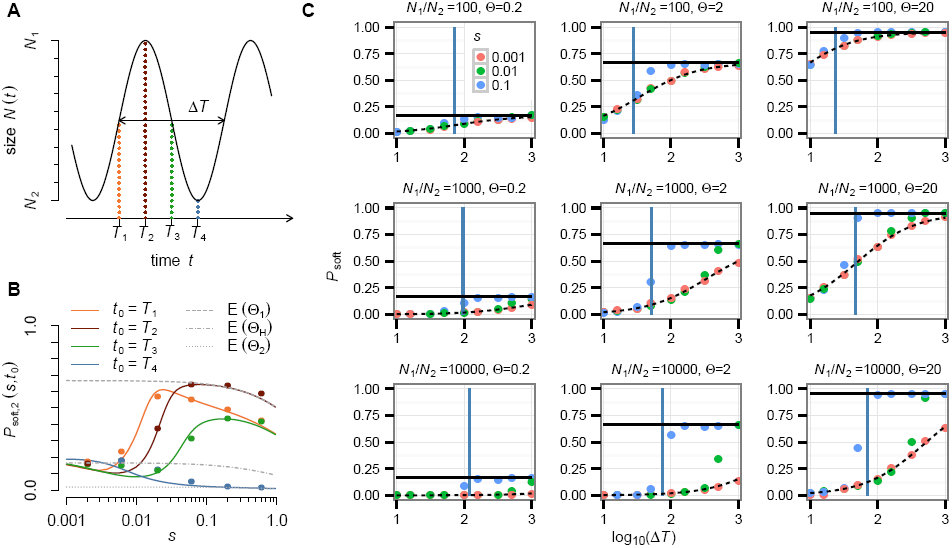
Weak and strong selection limits. (A) In the cyclical population model, *N*(*t*) cycles between a maximum size *N*_max_ = 10^8^ and a minimum size *N*_min_ = 10^6^ over a period of Δ*T* = 500 generations. Adaptive mutations occur at a *de novo* rate of *U*_*A*_ = 10^−8^ per individual, per generation. We condition selective sweeps on four different starting times: *T*_1_, *T*_2_, *T*_3_, and *T*_4_. (B) Comparison of our analytical predictions for the probabilities *P*_soft,2_(*t*) of observing a soft sweep in a sample of two adaptive alleles, drawn randomly at the time when the adaptive allele has reached a population frequency of 50% (colored lines), with empirical probabilities observed in Wright-Fisher simulations (colored circles, see Methods). Convergence to the harmonic mean expectation, *E*(Θ_*H*_), is seen for weak selection, while convergence to the instantaneous population size expectation, *E*(Θ_1_) and *E*(Θ_2_), is seen for strong selection. The convergence of the orange and light blue lines is also expected in the strong selection limit as they share the same instantaneous population size at *t*_0_. (C) The weak selection / fast fluctuation and strong selection / slow fluctuation limits are also observed in our recurrent bottleneck model from Figure 2. The observed probabilities of soft sweeps in the recurrent bottleneck simulations transition from the harmonic mean expectations (dashed black lines) to the instantaneous population size expectations (solid black lines). The solid blue vertical line indicates the position of our heuristic boundary 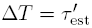 for the selection coefficient *s* = 0.1.

We derived our analytical predictions for *P*_soft,2_(*t*_0_, *s*) by first calculating *P*_fix_(*t*_0_, *s*) for the given *N*(*t*), *t*_0_, and *s* via numerical integration of Equation (9) and then inserting the result into Equation (8) to obtain the trajectory *x*\*(*t*), using the scaling *N*_*e*_(*t*) = *N*(*t*)/(1 + *s*) for concordance between the generalized birth-death model used by Uecker and Hermisson (2011) and the Wright-Fisher model. We then estimated *E*(*t*_1_) via numerical integration of Equation (10) (Methods), assuming that the adaptive allele reaches frequency *x* = 1/2 at:

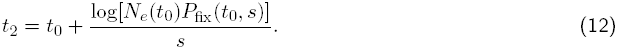

Figure 4B shows the comparison between our analytical predictions for *P*_soft,2_(*t*_0_, *s*) and the observed frequencies of soft sweeps in Wright-Fisher simulations for a scenario with population sizes *N*_min_ = 10^6^ = 0.01(*N*_max_), cycle period Δ*T* = 500, and adaptive mutation rate *U*_*A*_ = 10^−8^, as a function of the strength of positive selection and the starting time of the sweep within a cycle. Simulation results are in good agreement with analytical predictions over the whole range of investigated parameters.

We observe two characteristic limits in our cyclical population size model, specified by the relation between the duration of the sweeps (which inversely depends on the selection strength) and the timescale over which demographic processes occur:

i. **weak selection / fast fluctuation limit:** When the duration of a sweep becomes much longer than the period of population size fluctuations, the probability of observing a soft sweep converges to that expected in a population of constant size, given by the harmonic mean of *N*_*e*_(*t*) estimated over a population cycle (dash-dotted line in Figure 4B). The starting time of the sweep becomes irrelevant in this case. To show this, we partition the embedded integral *∫*1/(*N*_*e*_*x*)*dt* in Equation (6) into consecutive intervals, each extending over one population cycle. Because *x*(*t*) changes slow compared with the timescale of a population cycle, we can assume that *x*(*t*) is approximately constant over each such interval. The harmonic mean then factorizes into *H*_*k*_(*N*_*e*_*x*) = *H*_*k*_(*N*_*e*_)*H*_*k*_(*x*), and Equation (6) reduces to

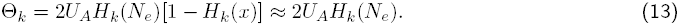 The last approximation holds as long as *k* is not too large, in which case the lowest value of *x*(*t*) in the interval, and thus also *H*_*k*_(*x*), are still small, since the harmonic mean is dominated by the smallest values. Note that the above argument applies more broadly and is not necessarily limited to scenarios where population size fluctuations are exactly cyclical. In general, a sufficient condition for the factorization in Equation (13) is the existence of a timescale, *ξ*, that is much shorter than the duration of the sweep, where harmonic averages of *N*(*t*) estimated over time intervals of length *ξ* are already approximately constant for every interval lying within the duration of the sweep. In other words, factorization works for all demographic models that have fast fluctuation modes we can effectively average out but no slow fluctuation modes occurring over timescales comparable to the duration of the sweep. Examples for demographic models where the weak selection / fast fluctuation limit becomes applicable include those where *N*(*t*) is any periodic function with a period much shorter than the duration of the sweep. Another example would be a model in which population sizes are drawn randomly from a distribution with fixed mean, where the number of drawings over the duration of the sweep is large enough such that harmonic averages already converge to the mean over timescales much shorter than the duration of the sweep.
ii. **strong selection / slow fluctuation limit:** When the duration of a sweep becomes much shorter than the timescale over which population size changes, the probability of observing a soft sweep in the cyclical population model converges to that which is expected in a population of constant size *N*_*e*_(*t*_0_), the effective population size at the starting time of the sweep. In this case the effective Θ_*k*_ from Equation (6) reduces to

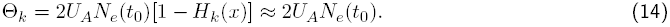 We can also recover these weak and strong selection limits for our earlier simulations of the recurrent bottleneck scenario. Figure 4C shows the transition from what is expected in a constant population given by the harmonic mean population size over one bottleneck cycle, *H*(*N*_*e*_), to a constant population at the instantaneous population size, *N*_*e*_(*t*_0_) = *N*_1_. The expectations in the limits were calculated using Equation (1) while substituting the appropriate effective population size. Again we see that even for the same demographic scenario, the probability of observing a soft sweep can vary dramatically with selection coefficient. This implies that there is generally no one effective population size that will be relevant for determining the expected selective sweep signature. Notice also that while the transition between the two regimes in our hardening model is monotonic, the transition is not guaranteed to be monotonic in more complex demographic scenarios, as seen for some of the transitions in our cycling population model.

## Discussion

In this study we investigated the population parameters that determine the probability of observing soft selective sweeps when adaptation arises from *de novo* mutations. Our understanding of soft sweeps has hitherto been limited to the special case where population size remains constant over time. In this special case, the probability of soft sweeps from recurrent *de novo* mutation depends primarily on the parameter Θ = 2*N*_*e*_*U*_*A*_ (twice the population-scale mutation rate towards the adaptive allele) and is largely independent of the strength of selection (Pennings and Hermisson 2006a). We devised a unified framework for calculating the probability of observing soft sweeps when population size changes over time and found that the strength of selection becomes a key factor for determining the likelihood of observing soft sweeps in many demographic scenarios.

**The hardening phenomenon:** We first demonstrated that population bottlenecks can give rise to a phenomenon that we term the hardening of soft selective sweeps. Hardening describes a situation where several adaptive mutations of independent origin – initially destined to produce a soft sweep in a constant population – establish in the population, but only one adaptive lineage ultimately survives a subsequent bottleneck, resulting in a hard selective sweep.

Using a simple heuristic approach that models the trajectories of adaptive alleles forward in time, we showed that in populations that experience recurrent, sharp bottlenecks, the likelihood of such hardening depends on the comparison of two characteristic timescales: (i) the recurrence time (Δ*T*) between bottlenecks and (ii) the bottleneck establishment time (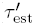) which specifies the waiting time until a *de novo* adaptive mutation reaches a high-enough frequency such that it is virtually guaranteed to survive a bottleneck. We derived a simple heuristic approximation, 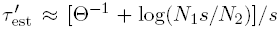, that applies when bottlenecks are severe enough (*N*_1_*s* > *N*_2_ and *N*_2_ ≫ 1). If soft sweeps are expected to arise between bottlenecks – i.e., if Θ is on the order of one or larger during those phases – then hardening is common when 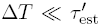, whereas it is unlikely when 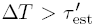. The bottleneck establishment time increases only logarithmically with the severity of the bottleneck and scales inversely with the selection coefficient of the adaptive mutation. In stark contrast to a population of constant size, the probability of observing soft sweeps can therefore strongly depend on the strength of selection in the recurrent bottleneck scenario.

**Generalized analytical framework for complex demographies:** The heuristic condition 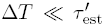 provides a rough estimate of whether hardening is expected in a recurrent bottleneck scenario, but it lacks generality for more complex demographic scenarios and does not provide the actual probabilities of observing soft sweeps. We showed that such probabilities can be calculated analytically for a wide range of demographic models by mapping the problem onto a coalescent with killings process (Durrett 2008). Our approach is very similar to that employed by Pennings and Hermisson (2006a) for the constant size model, with the primary difference being that we allow for coalescence and mutation rates to vary over time as population size changes.

In the coalescent with killings framework (Figure 3), the probability of a soft sweep is determined by the competition between two processes: coalescence of adaptive lineages in the fraction *x*(*t*) of the population that carry the adaptive allele, and emergence of new adaptive lineages through *de novo* mutation (referred to as killings when going backwards in time) in the fraction 1 − *x*(*t*) of the population that do not yet carry the adaptive allele. A sweep is hard in a population sample if all lineages in that sample coalesce before a second adaptive mutation arises and soft otherwise. In our analytical approach, we assume that the trajectory *x*(*t*) can be described by a logistic function. The probability of observing a soft sweep can then be calculated through numerical integration of the expected rates of coalescence and mutation in the genealogy, which are simple functions of *x*(*t*) and *N*_*e*_(*t*), the variance effective population size in generation *t*.

Note that by adjusting the end-point of the integration interval to the time at which the adaptive allele reaches a given frequency, our approach can easily be extended to the analysis of partial selective sweeps. Similarly, by extending the time interval beyond the fixation of the adaptive allele, one can study the loss of adaptive lineages due to random genetic drift after the completion of a soft sweep. Moreover, since our model only requires an estimate of the frequency trajectory of the adaptive allele, *x*(*t*), it should be easily extendable to other, more complex scenarios, including time-varying selection coefficients (Uecker and Hermisson 2011), as long as one can still model *x*(*t*) in the particular scenario. We leave these possible extensions for future exploration.

Even though the results presented in this paper were derived for haploid populations, it is straightforward to extend them to other levels of ploidy. The key prerequisite is again that we still have an estimate for the frequency trajectory of the adaptive allele, which can be complicated by dominance effects when ploidy increases. Given the trajectory, the population size *N*(*t*) simply needs to be multiplied by the ploidy level to adjust for the changed rate of coalescence in the genealogy. For example, in a diploid population with codominance, the population-scale mutation rate needs to be defined as Θ = 4*N*_*e*_*U*_*A*_, twice the value for a haploid population of the same size.

**Weak and strong selection limits:** Our approach reveals interesting analogies to Kingman’s coales-cent (Kingman 1982) with respect to our ability to map the dynamics onto an effective model of constant population size. SjÖdin *et al.* (2005) showed that genealogies at neutral loci can be described by a linear rescaling of Kingman’s coalescent with a corresponding coalescent effective population size, as long as demographic processes and coalescence events operate on very different timescales. Specifically, when population size fluctuations occur much faster compared with the timescale of coalescence, the coalescent effective population size is given by the harmonic mean of the variance effective population size, *N*_*e*_(*t*), estimated over the timescale of coalescence. In the opposite limit where population size fluctuations occur much slower than the timescales of neutral coalescence, the variance effective population size is approximately constant over the relevant time interval and directly corresponds to the instantaneous coalescent effective population size.

Analogously, in our analytical framework for determining the likelihood of soft sweeps, we can again map demography onto an effective model with constant effective population size in the two limits where population size fluctuations are either very fast or very slow. The relevant timescale for comparison here is the duration of the selective sweep, *τ*_fix_ ≈ log(*N*_*s*_)/*s*, which is inversely proportional to the selection coefficient of the sweep. Hence, the fast fluctuation limit corresponds to a weak selection limit, and the slow fluctuation limit to a strong selection limit. In the strong selection / slow fluctuation limit, the relevant effective population size is the instantaneous effective population size at the start of the sweep; in the weak selection / fast fluctuation limit, it is the harmonic mean of the variance effective population size estimated over the duration of the sweep.

One important consequence of this finding is that, even in the same demographic scenario, the probability of observing soft sweeps can differ substantially for weakly and strongly selected alleles. This is because the harmonic mean that determines the effective population size in the weak selection / fast fluctuation limit will be dominated by the phases where population size is small. For a weakly selected allele in a population that fluctuates much faster than the duration of the sweep, it will be close to the minimum size encountered during the sweep, resulting in a low effective population size and, correspondingly, a low probability of observing a soft sweep. A strongly selected allele, on the other hand, can arise and sweep to fixation between collapses of the population. The effective population size remains large in this case, increasing the probability of observing a soft sweep. Hence, the stronger the selective sweep, the higher the chance that it will be soft in a population that fluctuates in size.

Similar behavior is observed for the fixation probabilities of adaptive alleles in fluctuating populations. In particular, Otto and Whitlock (1997) showed that the fixation process of an adaptive allele depends on the timescale of the fixation itself. Only short-term demographic changes encountered during the fixation event matter for strongly selected alleles, whereas slower changes only affect weakly selected alleles. Otto and Whitlock (1997) therefore concluded that “there is no single effective population size that can be used to determine the probability of fixation for all new beneficial mutations in a population of changing size.”

**Hard versus soft selective sweeps in natural populations:** How relevant is our finding that the likelihood of observing soft sweeps can strongly depend on the strength of selection for understanding adaptation in realistic populations? We know that both necessary ingredients for this effect to occur – strong temporal fluctuations in population size and difference in the fitness effects of *de novo* adaptive mutations – are common in nature.

Population size fluctuations over several orders of magnitude are observed in various animal species, ranging from parasitic worms to insects and even small mammals (Berryman 2002). Unicellular organisms often undergo even more dramatic changes in population size. For instance, during Malaria infection only ten to a hundred sporozoites are typically ejected by a feeding mosquito – the numbers of sporozoites that successfully enter the human blood stream are even smaller – yet this population grows to many billions of parasites within an infected individual (Rosenberg *et al.* 1990). Similarly, in the majority of cases acute HIV infection was found to result from a single virus (Keele *et al.* 2008). Severe population bottlenecks resulting from serial dilution are also commonly encountered in evolution experiments with bacteria and yeast (Wahl *et al.* 2002). Even our own species has likely experienced population size changes over more than three orders of magnitude within the last 1000 generations (Gazave *et al.* 2014).

It is also well established that fitness effects of *de novo* adaptive mutations can vary over many orders of magnitude within the same species. For example, codon bias is typically associated with only weak selective advantages, whereas the fitness advantage during the evolution of drug resistance in pathogens or pesticide resistance in insects can be on the order of 10% or larger.

Taken together, we predict that we should be able to observe strong dependence of the likelihood of hardening on the strength of selection for adaptation in natural populations that experience a demographic phase where adaptation is not mutation-limited. The likelihood of observing soft sweeps will depend on the types of natural population fluctuations that occur and whether they can be characterized by the weak selection / fast fluctuation limit or the strong selection / slow fluctuation limit.

To demonstrate this possibility, consider a cycling population illustrated in Figure 5A that is based on data from the extreme fluctuations observed in multiple species of moths, including the tea tortrix, *Adoxophyes honmai*, and the larch budmoth, *Zeiraphera diniana*. These diploid moth species have been observed to undergo changes in population size spanning many orders of magnitude over short periods of just four to five generations (Baltensweiler and Fischlin 1988; Nelson *et al.* 2013). Let us further assume that these changes result in a change in the adaptive population-scale mutation rate between Θ_min_ = 10^−3^ and Θ_max_ = 1. In this case, adaptation is not mutation-limited during population maxima and is mutation-limited during population minima. Consequently, hardening of soft selective sweeps is expected to be common.

**Figure 5:**
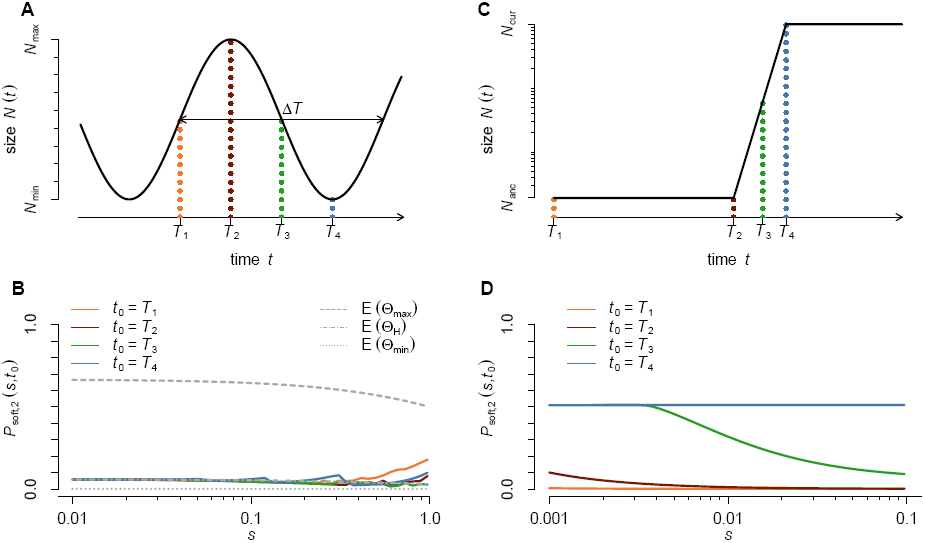
Probability of observing soft sweeps in two demographic scenarios. (A) Fluctuating population inspired by data from the extreme fluctuations observed in multiple species of moths (Baltensweiler and Fischlin 1988; Nelson *et al.* 2013). We assume the adaptive population-scale mutation rate varies between Θ_min_ = 10^−3^ and Θ_max_ = 1 over a period of Δ*T* = 5 generations. (B) Our predictions for the probability of observing a soft sweep in a sample of two adaptive alleles drawn randomly at the time-point when the adaptive allele has reached a population frequency of 50%, conditional on four different starting times of the sweep (*T*_1_ to *T*_4_). The noise in the predictions stems from the numerical Monte Carlo integrations in our approach. The probability of observing a soft sweep is close to the harmonic mean expectation, *E*(Θ_*H*_), for virtually all starting times and selection strengths, except when selection is extremely strong. (C) Demographic model proposed for the European human population (Gazave *et al.* 2014). The ancestral population size is *N*_anc_ = 10^4^. Starting at 113 generations in the past, the population expands exponentially at a constant rate of *r* = 0.0554, until it reaches its current size of *N*_cur_ ≈ 520,000. Population size is assumed to remain constant thereafter. Note that the y-axis is plotted logarithmically. We set the beneficial mutation rate in this example at *U*_*A*_ = 5 × 10^−8^. (D) Analytic predictions for the probability of observing a soft sweep in a sample of size two when the sweep starts at present (*T*_4_), midway during the expansion (*T*_3_ = 50 generations ago), at the beginning of the expansion (*T*_2_ = 113 generations ago), and prior to the expansion (*T*_1_ = 500 generations ago). Sweeps that start prior to the expansion are almost exclusively hard, whereas sweeps starting today will be soft in more than half of the cases, regardless of the strength of selection. Sweeps starting at the beginning or during the expansion show an interesting crossover behavior: smaller selection coefficients are more likely than larger selection coefficients to produce soft sweeps because weaker sweeps take longer to complete and thus experience more time at larger population sizes.

Figure 5B shows the likelihood of soft sweeps in this scenario according to Equation (7), as a function of the strength of selection and the starting time of the sweep. The probability of observing soft sweeps generally remains low in this scenario, except for cases of extremely strong selection. We can understand this result from the fact that the timescale of population size fluctuations is so fast that all but the most strongly selected alleles still fall within the weak selection limit, described by the harmonic mean.

This result has important consequences for the study of other populations that fluctuate over similarly short timescales, such as the fruit fly *Drosophila melanogaster*. Natural populations of *Drosophila melanogaster* undergo approximately 10–20 generations over a seasonal cycle, often experiencing enormous population sizes during the summer that collapse again each winter (Ives 1970). Our result then suggests that only the most strongly selected alleles, which can arise and sweep over a single season, may actually fall within the strong selection limit. All other sweeps should still be governed by the harmonic mean of the population size averaged over a yearly cycle, which will be dominated by the small winter population sizes.

Let us consider another example, motivated by the proposed recent demographic history of the European human population (Coventry *et al.* 2010; Nelson *et al.* 2012; Tennessen *et al.* 2012; Gazave *et al*. 2014). Specifically, we consider a population that was small throughout most of its history and has recently experienced a dramatic population expansion. We assume demographic parameters similar to those estimated by Gazave *et al.* (2014), *i.e.*, an ancestral population size of *N*_anc_ = 10^4^, followed by exponential growth over a period of 113 generations, reaching a current size of approximately *N*_cur_ ≈ 520,000 individuals (Figure 5C). We further assume that exponential growth halts at present and that population size remains constant thereafter. Note that this scenario is qualitatively different from the previously discussed models in that population size changes are non-recurring. As a result, the weak selection / fast fluctuation does not exist in this case. For determining whether a given selective sweep will likely be hard or soft in this model, its starting time becomes of crucial importance.

We assume an adaptive mutation rate of *U*_*A*_ = 5 × 10^−7^ for this example to illustrate the transition between mutation-limited behavior in the ancestral population, where Θ_anc_ = 4*N*_anc_*U*_*A*_ ≈ 0.02, and non-mutation-limited behavior in the current population, where Θ_cur_ = 4*N*_cur_*U*_*A*_ ≈ 1.0. Note that this adaptive mutation rate is higher than the single nucleotide mutation rate in humans, but it may be appropriate for describing adaptations that have larger mutational target size, such as loss-of-function mutations or changes in the expression level of a gene. Moreover, if we were to assume that the current effective population size of the European human population is in fact *N*_cur_ ≈ 2 × 10^7^ – still over an order of magnitude smaller than its census size – we would already be in the non-mutation-limited regime for *U*_*A*_ ≈ 10^−8^, the current estimate of the single nucleotide mutation rate in humans (Kong *et al.* 2012).

Figure 5D shows the probabilities of soft sweeps in this scenario predicted by our approach as a function of the strength of selection and starting time of the sweep. The results confirm our intuition that almost all sweeps that start prior to the expansion are hard in a sample of size two, as expected for adaptation by *de novo* mutation in a mutation-limited scenario, whereas sweeps starting in the current, non-mutation-limited regime are soft in more than half of the cases, regardless of the strength of selection. Sweeps starting during the expansion phase show an interesting crossover behavior between hard and soft sweeps. The strength of selection becomes important in this case. Specifically, sweeps that start during the expansion have a higher probability of producing soft sweeps when they are driven by weak selection than when they are driven by strong selection. This effect can be understood from the fact that stronger sweeps go to fixation faster than the weaker sweeps. Hence, in a growing population, a weak sweep will experience larger population sizes during its course than a strong sweep starting at the same time, increasing its probability of beeing soft.

When expanding the intuition from our single-locus model to whole genomes, we must bear in mind that the effective Θ determining the probability of soft sweeps will not be the same for different loci across the genome because mutation rate and target size will vary for adaptive mutations at different loci. For example, adaptive loss-of-function mutations will likely have a much higher value of *U*_*A*_ than adaptive single nucleotide mutations. Therefore, no single value of Θ will be appropriate for describing the entire adaptive dynamics of a population. Adaptation across the genome can simultaneously be mutation-limited and non-mutation-limited in the same population, depending on population size fluctuations, mutation rate, target size, and the strength of selection. Furthermore, we should be very cautious when assuming that estimators for Θ based on genetic diversity will inform us about whether recent adaptation will produce hard or soft sweeps. Estimators based on the levels of neutral diversity in a population, such as Θ_*π*_ and Watterson’s Θ_*W*_ (Ewens 2004), can be strongly biased downward by ancient bottlenecks and recurrent linked selection.

Finally, the overall prevalence of soft sweeps should depend on when adaptation and directional selection is common. If adaptation is limited by mutational input, then most adaptive mutations should arise during the population booms, biasing us toward seeing more soft sweeps. On the other hand, it is also possible – maybe even more probable – that adaptation will be common during periods of population decline, such as when population decline is caused by a strong selective agent like a new pathogen, competitor, predator, or a shortage in the abundance of food. If adaptation is more common during population busts, this should lead us to observe more hard sweeps.

These considerations highlight one of the key limits of the current analysis – we have only considered scenarios where population size and selection coefficients are independent of each other. In the future, we believe that models that consider population size and fitness in a unified framework will be necessary to fully understand signatures that adaptation leaves in populations of variable size.

## Methods

**Forward simulations of adaptation under recurrent population bottlenecks:** We simulated adaptation from *de novo* mutation in a modified Wright-Fisher model with selection. Each simulation run was started from a population that was initially monomorphic for the wildtype allele, *a*. New adaptive mutations entered the population by a Poisson process with rate *N*_1_*U*_*A*_[1 − *x*(*t*)], where 1 − *x*(*t*) is the frequency of the wildtype allele. The population in each generation was produced by multinomial sampling from the previous generation, with sampling probabilities being proportional to the difference in fitness of each lineage and the mean population fitness. Population bottlenecks were simulated through a single-generation downsampling to size *N*_2_ (without selection) every Δ*T* generations. We did not require that the first beneficial mutation arise in the first generation. Each simulation run started Δ*T* generations before the first bottleneck. All adaptive lineages were tracked in the population until the adaptive allele had reached fixation. One thousand simulations were run for each parameter combination. Empirical probabilities of observing a soft sweep in a given simulation run were obtained by calculating the expected probability that two randomly drawn adaptive lineages are not identical by decent, based on the population frequencies of all adaptive lineages in the population at the time of sampling. All code was written in Python and C++ and is available upon request.

**Numerical Monte Carlo integration:** Analytical predictions for *P*_soft,2_(*t*, *s*) in Figure 4 and Figure 5 were obtained by the following procedure: For the given demographic model, selection coefficient, and starting time of the sweep, we first calculated the fixation probability of the adaptive allele via Equation (9) using Monte Carlo integration routines from the GNU Scientific Library (Galassi *et al.* 2009). This fixation probability was then used in Equation (8) to obtain the deterministic trajectory *x*\*(*t*). Solving *x*\*(*t*_2_) = 1/2 yielded the sampling time *t*_2_. We then iteratively estimated the lower bound *E*(*t*_1_) of integral (10) such that the expected number of events occurring between *E*(*t*_1_) and *t*_2_ converged to 1±10^−4^. Finally, we integrated the coalescence rate from Equation (4) over the interval [*E*(*t*_1_), *t*_2_] to determine the probability that the event occurring at *E*(*t*_1_) was a coalescent event, yielding *P*_coal_ = 1 − *P*_soft,2_. Note that this approach can easily be adjusted for any other sampling time or adaptive allele frequency at sampling.

**Forward simulations in cycling populations:** We simulated adaptation from *de novo* mutation in a cycling population using the Wright-Fisher model specified above. Each simulated population was initially monomorphic for the wildtype allele. We began our simulations at four different time points (*t*_0_) along the population growth cycle and ran each simulation on the condition that the first beneficial allele that arose in generation *t*_0_ did not go extinct during the simulation. Simulations were run until the adaptive allele was above 50% frequency. Ten thousand simulations were run for each combination of parameters.

## Acknowledgments

We thank Pleuni Pennings, Jamie Blundell, and Hildegard Uecker for useful discussions leading to the formulation of our primary results. We thank Nandita Garud, Joachim Hermisson, Marc Feldman, Daniel Fisher, and members of the Petrov lab for comments and suggestions made prior to and during the formulation of this manuscript. B.A.W. is supported by the NSF Graduate Research Fellowship. This work was supported by the NIH under grants GM089926 and HG002568 to D.A.P.

